# DLC3/Cv-c function in testis development in humans and Drosophila: implication for variants of sex development

**DOI:** 10.1101/2022.07.28.501838

**Authors:** Sol Sotillos, Isabel von der Decken, Ivan Domenech Mercadé, Sriraksha Srinivasan, Stefano Vanni, Serge Nef, Anna Biason-Lauber, Daniel Rodríguez Gutiérrez, James C-G Hombría

## Abstract

Identifying genes affecting gonad development is essential to understand the mechanisms causing Variants/Differences in Sex Development. Recently, a *DLC3* mutation was associated with male gonadal dysgenesis in 46,XY DSD patients. We show that Cv-c, the *Drosophila* ortholog of DLC3, is also required to maintain testis integrity during fly development. We found that Cv-c and human DLC3 can perform the same function in fly embryos, as flies with wild type but not mutated DLC3 rescue gonadal dysgenesis, suggesting a functional conservation. Expression of different Cv-c protein variants demonstrated that the StART domain mediates the Cv-c function in the male gonad, independently from the GAP domain activity. This work demonstrates a role for DLC3/Cv-c in male gonadogenesis and highlights a novel StART-mediated function required for gonadal mesoderm-germ cell interaction during testis development.

**Significance Statement:** Associating rare human genetic variants to specific conditions is complex. An amino acid change in the StART domain of the RhoGAP DLC3 protein has been found in female DSD patients with a 46,XY male karyotype. We present a second DSD patient with a mutation in the same region and show that the Cv-c/DLC3 homolog is also required in Drosophila testis. In *cv-c* mutant embryos the testis mesodermal cells display various defects: the testes are unable to retain the germ cells due to their abnormal ensheathment by mesodermal cells and the mesodermal sheet of cells surrounding the testis is discontinuous resulting in the liberation of the germ cells outside the gonad. Defects can be rescued by gonadal expression of Cv-c or DLC3 but not by the patients’ DLC3 protein variant. Testis development requires the StART lipid binding domain but not the GAP domain, revealing a novel function of this RhoGAP family.

## Introduction

Human sex development relies on the correct differentiation and function of the gonads. This requires a delicate functional balance between genes, cells and hormones. Mutations affecting the determination and differentiation of the gonads can lead to Differences of Sex Development (DSD) where the female (XX) or male (XY) sex karyotype does not match with gonadal and anatomical development (Cools *et al*, 2018). The primary cause of nearly 50% of DSD cases remains unknown (Barseghyan *et al*, 2018; Buonocore *et al*, 2019), suggesting the existence of further sex-developmental mechanisms still awaiting discovery.

Gonadogenesis can be subdivided into three stages: specification of precursor germ cells, directional migration towards the somatic gonadal precursors and gonad compaction. In mammals, somatic cells, i.e. Sertoli cells in male and Granulosa cells in females, play a central role in sex determination with the germ cells differentiating into sperms or oocytes depending on their somatic mesoderm environment. In humans, Primordial Germ Cells (PGCs) are formed near the allantois during gastrulation around the 4^th^ gestational week (GW) and migrate to the genital ridge where they form the anlage necessary for gonadal development (GW5-6). Somatic mesodermal cells are required for both PGCs migration and the formation of a proper gonad. Once PGCs reach their destination, the somatic gonadal cells join them (around GW 7-8 in males, GW10 in females) and provide a suitable environment for survival and self-renewal until gamete differentiation (Jemc, 2011). Thus, mutations in genes regulating somatic Sertoli and Granulosa support cell function in humans are often associated with complete or partial gonadal dysgenesis in both sexes and sex reversal in males (Brunello & Rey, 2021; Knower *et al*, 2011; Zarkower & Murphy, 2021). Other mesodermal cells, the Leydig cells, also play an important role in the testis by being the primary source of testosterone and other androgens and maintaining secondary sexual characteristics.

The central elements of gonadogenesis are relatively well conserved among species. In *Drosophila*, PGCs are formed in the blastoderm and carried passively into the gut where they enter the embryo after crossing the intestinal epithelium. PGCs migrate towards their final position where they coalesce forming a compact gonad. The dependency of PGCs on somatic gonadal cells during development is also well conserved. In mouse mutants without a genital ridge, the PGCs can migrate but remain immature (reviewed in (Cooke & Moris, 2021)). In the fly, somatic gonadal cells can coalesce into a gonad in the absence of PGCs, but the PGCs are unable to coalesce in the absence of somatic cells (Brookman *et al*, 1992). Similarly, subsets of somatic gonadal cells produce steroid hormones (testosterone in mammals (Zirkin & Papadopoulos, 2018) and ecdysone in insects (Tajouri *et al*, 2018)), which have a global influence on the organism.

Abnormal gonadogenesis in human 46,XY individuals leads to under-masculinization, resulting in incomplete sexual characteristics at birth. Patients with complete gonadal dysgenesis present female external genitalia and hypogonadotropic hypogonadism with lack of secondary sex characteristics (Rocha *et al*, 2011). In mammalian models, gonadal dysgenesis causes infertility, DSD and sex reversal while in insect models it leads to sterility.

Recently, the human X-linked *DLC3* human gene (also known as *STARD8*) has been implicated in a case of 46,XY gonadal dysgenesis in two patients carrying a variant in the StART domain (Ilaslan *et al*, 2018). The Deleted in Liver Cancer (DLC) proteins belong to the RhoGAP family of small GTPase regulators. In vertebrates there are three members (DLC1,2 and 3) whereas *Drosophila* has a single ortholog, *crossveinless-c* (*cv-c*) (Denholm *et al*, 2005). This family of proteins share different domains: besides the Rho <u>G</u>TPase <u>A</u>ctivating <u>P</u>rotein domain (GAP), they present a protein-protein interacting <u>S</u>terile <u>A</u>lpha <u>M</u>otif (SAM) at the N terminal end and a <u>St</u>eroidogenic <u>A</u>cute <u>R</u>egulatory protein (StAR)-related lipid transfer (StART) domain at the C terminal. StART domains have been shown in other proteins to be involved in lipid interaction, protein localization and function (Braun & Olayioye, 2015; Clark, 2020).

We previously reported that DLC1 and DLC3 can functionally substitute for Cv-c in *Drosophila* (Sotillos *et al*, 2018) opening up the use of *Drosophila* as a system to analyse the requirement of DLC3/Cv-c proteins during male gonadogenesis. Here, using *Drosophila,* we demonstrate that the RhoGAP Cv-c and DLC3 proteins have a conserved role in male gonad formation mediated by the StART domain, confirming the suspected DLC3 involvement in human testicular organogenesis.

## Results

### The S993N mutation is located in a functionally-important region of the DLC3-StART domain and can alter its conformational dynamics

The extremely rare (MAF=0.00016 according to Gnomad) *DLC3* mutation was previously observed in two related DSD patients where a Serine (S) amino acid (aa) was substituted by an Asparagine (N) at position 993. The mutation was present in the heterozygous carrier mother (Fig. 1*A*) and in the two 46,XY dysgenic patients but not in their 46,XY healthy sibling (Ilaslan *et al*., 2018). However, despite this strong correlation, no direct evidence linking DLC3 to gonadal dysgenesis was provided.

**Figure 1:**
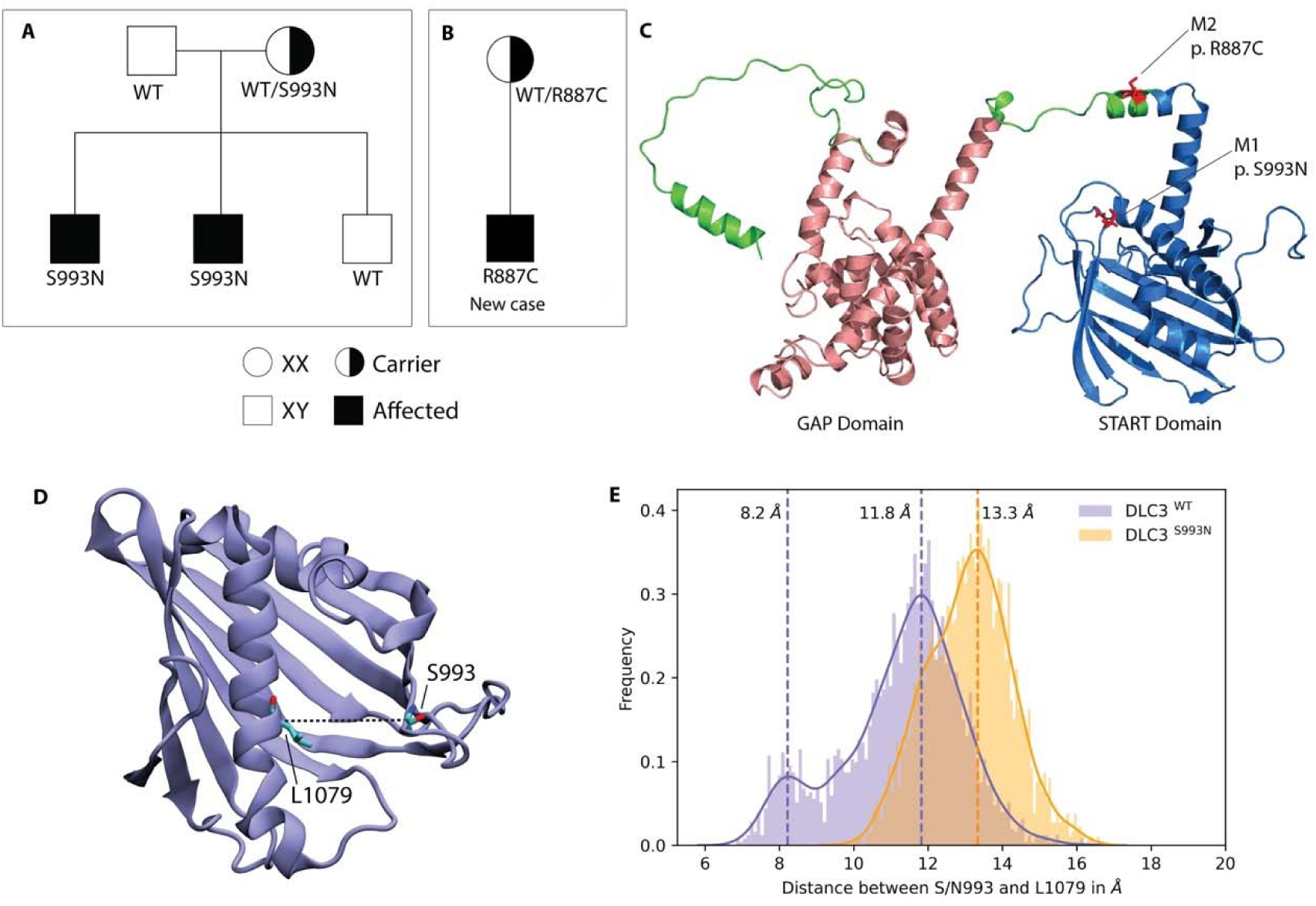
DLC3 variants associated to DSD patients. (**A**-**B**) Family diagrams showing the segregation of two different DLC3 alleles (panel A is modified from Ilaslan, E. et al. 2018). (**C**) Structure of DLC3 GAP and StART domains showing the localization of the pR887C and the pS993N mutations. The protein is shown in cartoon representation, with the GAP domain represented in orange and the StART domain in blue. (**D**) Structure of the StART domain of human DLC3/StARD8. The distance between the Cα atoms of residues S/N993 and L1079 (shown in licorice representation) were used to determine the open and closed transitions arising from the motion of the Ω1 loop (**E**) Distribution of the distance between the Cα atoms of S/N993 and L1079 in atomistic simulations of the WT and S993N systems.

Recently, we have identified a third DSD patient carrying a DLC3 variant, who inherited the R887C extremely rare mutation (MAF=0.00025) from the heterozygous mother (Fig. 1*B*). The patient presented 46,XY gonadal dysgenesis, where both gonads were not observable by ultrasonography. As this discovery reinforces the suspected involvement of DLC3 as a DSD factor, we decided to search for experimental evidence.

As a first approach we concentrated on the DLC3^S993N^ variant, which localizes to the conserved StART domain, analysing *in silico* if the aa substitution modifies the protein structure. The AlphaFold (AF) model of DLC3 indicates that residue S993 is located in the Ω1 loop of the StART domain (Fig. 1*C*), a loop that has been shown to be functionally important for several StART domains (Gatta *et al*, 2018; Horenkamp *et al*, 2018; Iaea *et al*, 2015; Khelashvili *et al*, 2019; Naito *et al*, 2019). To analyze if the S993N mutation affects the conformational dynamics of DLC3, we performed atomistic simulations of wild type DLC3 and of the mutant protein DLC3-S993N in water. We observed that the mutation causes a non-negligible effect on the conformation of the Ω1 loop that is located at the entrance of the hydrophobic cavity of the StART domain (Fig. 1*D* and *E*). The simulations of the wild type DLC3-StART domain show a transition between a “closed” state, where the distance between the Cα atoms of S993 and L1079 located on the opposite C-terminal helix is 8.3 Å, and an “open” state with an aperture of 11.9 Å. In contrast, the DLC3-S993N-StART domain showed a single “open” conformational state where the binding pocket has an aperture of 13.3 Å.

We also performed molecular dynamics (MD) simulations at the coarse-grain (CG) resolution level to predict membrane-interacting regions of the DLC3 StART domain (Supplementary Fig. 1) following a computational protocol we recently developed (Srinivasan *et al*, 2021). We observed multiple binding and unbinding events between the protein and the bilayer in all replicas, as indicated by the minimum distance between them (Supplementary Fig. 1A and B). When the frequency of interaction for each residue of the protein with the bilayer was determined (Supplementary Fig. 1C), the N-terminus of the protein and the Ω1 loop (Supplementary Fig. 1D) show the highest frequency of interaction.

These results indicate an effect of the S993N mutation on the conformational dynamics of the DLC3-StART domain and suggest that alterations of this Ω1 loop could impair the domain’s membrane interaction.

### Expression of the *cv-c RhoGAP* gene in the *Drosophila* gonads

Structural analyses have shown that DLC3 and Cv-c are highly conserved (Fig. 2*A* and *B*) and that, in *Drosophila*, DLC3 can functionally substitute for Cv-c (15). However, previous studies of Cv-c have concentrated in ectodermal derived tissues despite *cv-c* being broadly expressed in the mesoderm (Denholm *et al*., 2005; Sotillos *et al*, 2013; Sotillos *et al*., 2018). To find out if *cv-c* RNA is specifically transcribed in the gonadal mesoderm we performed fluorescent RNA *in situ* hybridization in embryos double stained with antibodies to detect gonad specific antigens. We observed that *cv-c* is transcribed in the testis mesoderm cells including the Somatic Gonadal Cells ensheathing the germ cells, the male-specific Somatic Gonad Precursors (msSGP) located at the posterior of the gonad and the Pigment Cell Precursors surrounding the whole testis (Fig. 2*C* and *D*). We confirmed Cv-c translation in these cells analysing the expression of a Cv-c::GFP fusion protein expressed under the endogenous *cv-c* regulatory elements (Fig. 2*F* and *H*). This Cv-c::GFP fusion protein has been shown to have identical distribution to the RNA expression (Sotillos *et al*., 2018). We did not observe comparable levels of *cv-c* mRNA nor Cv-c::GFP protein in the female gonad mesoderm (Fig. 2*E, G* and *I*) nor in the germ cells of any sex, indicating a male specific regulation of *cv-c* expression in the testis mesoderm.

**Figure 2:**
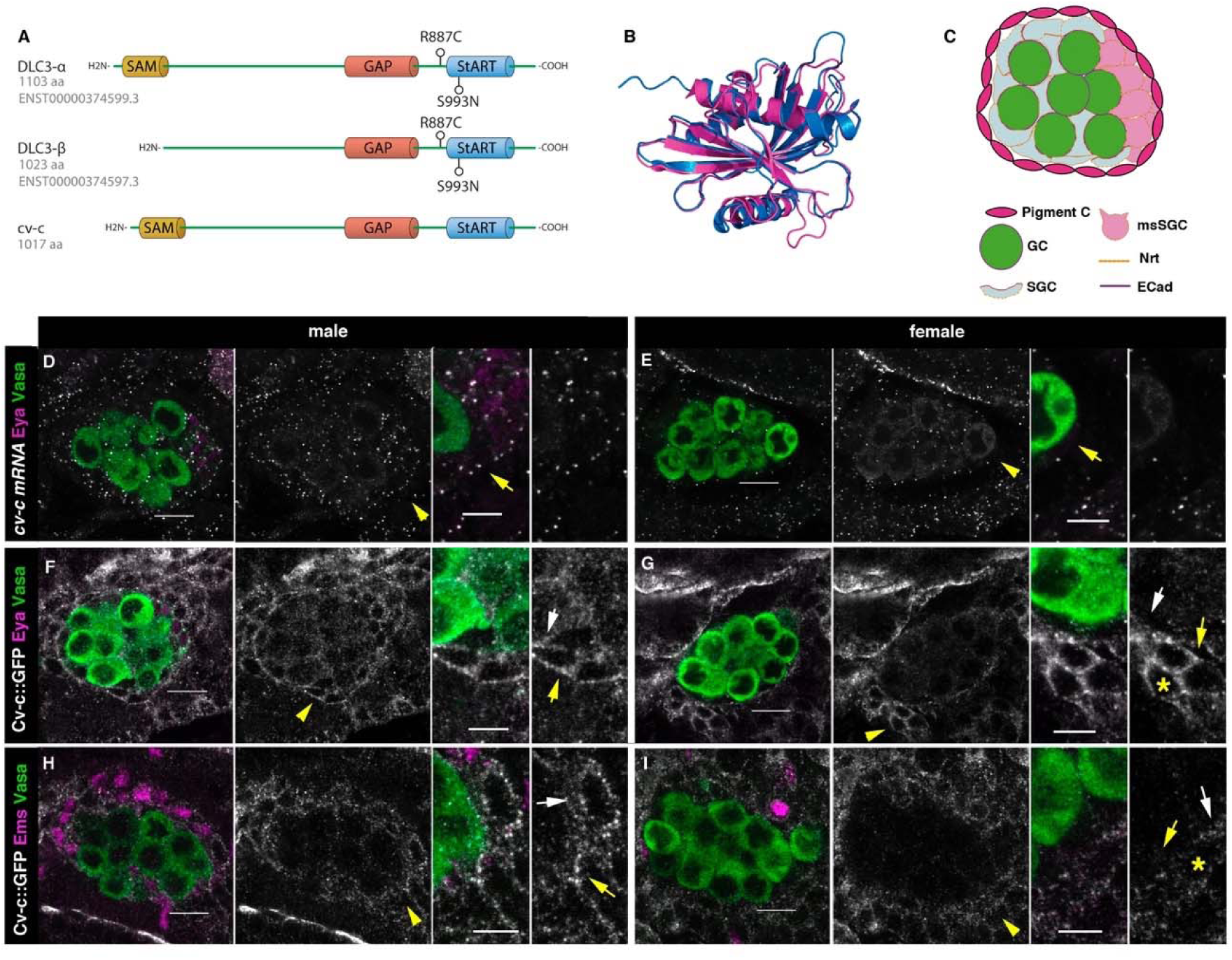
Cv-c and DLC3 structure and *cv-c* expression in the *Drosophila* gonad mesoderm. (**A**) Linear representation of the DLC3-α, DLC3-β, and Cv-c protein domains. The SAM domain is represented in yellow, the GAP domain in orange, and the StART domain in blue. (**B**) Alignment of the DLC3 (blue) and Cv-c (magenta) StART domains. (**C**) Schematic representation of the cell types in a *Drosophila* testis at st17. Germ Cells, green; Pigment Cells, magenta; Somatic Gonadal Cells, grey; male specific Somatic Gonadal Cells, pink. (**D**, **E**) RNA *in situ* hybridization of male (D) and female (E) st17 embryos shows general transcription of *cv-c* in the mesoderm. The right panels in D to I are close ups of the arrowed region in the central panels. (D) In the testis, comparable levels of mRNA puncta can be detected in the somatic mesoderm and in the gonadal mesoderm cells surrounding the male germ cells as is clearly observed in the msSGPs marked by Eya (magenta, indicated by an arrowhead in grey panels and an arrow in the close up). (E) In the ovary, marginal levels of *cv-c* mRNA expression are observed in the gonadal mesodermal cells, creating a halo of decreased number of puncta surrounding the female germ cells contrasting with the *cv-c* expressing adjacent somatic mesodermal cells (arrow in close up). (F-I) Cv-c::GFP protein expression in male and female embryos. (**F**, **H**) In the testis Cv-c::GFP is detected in the gonadal mesoderm surrounding the germ cells including the male msSGPs (Eya, magenta F) and the pigment cell precursors (Ems, magenta H). (**G**, **I**) In females, no substantial GFP signal is detected in the gonadal mesoderm surrounding the germ cells. Note in (F and H) that Cv-c::GFP signal in the gonad mesoderm cells allows tracing the testis contour, while in ovaries shown (G and I) this is not possible. Higher levels of Cv-c::GFP are present in the ectodermally derived trachea and hindgut. In close ups white arrows point to membranes close to the germ cells, yellow arrows to the membrane of gonad mesodermal cells. In males, Cv-c::GFP can be detected in the membranes between gonadal and somatic mesodermal cells (F, H) whilst in females GFP can only be detected outside the ovary in the membrane of the somatic mesoderm (G, I asterisks). Scale bar 10 µm and 5 µm in close ups.

### Cv-c functional requirement during male gonadogenesis in *Drosophila*

To test the functional significance of Cv-c expression in the fly gonads we analysed embryos homozygous for the lethal nonsense *cv-c^M62^*and *cv-c^C524^* alleles where stop codons result in truncated proteins lacking the GAP and StART domains (Denholm *et al*., 2005). We did not detect any major morphological defects in female gonads, confirming *cv-c* is not required for embryonic ovary development (Fig. 3*B*). In contrast, homozygous or hemizygous *cv-c* mutant male embryos have abnormal testes containing germ cells that are not surrounded by the gonadal mesoderm (Fig. 3*C* and *E* arrowheads compare with *A* and *D* respectively). This defect is unlikely to be due to the abnormal specification of the gonad mesoderm cells, as using mesodermal specific markers we can observe the presence of all cell types (Fig. 3*C, E* and *G*). However, the gonad mesoderm cells are frequently displaced, with the pigment cells failing to completely surround the mutant testis (Fig. 3*E*).

**Figure 3.**
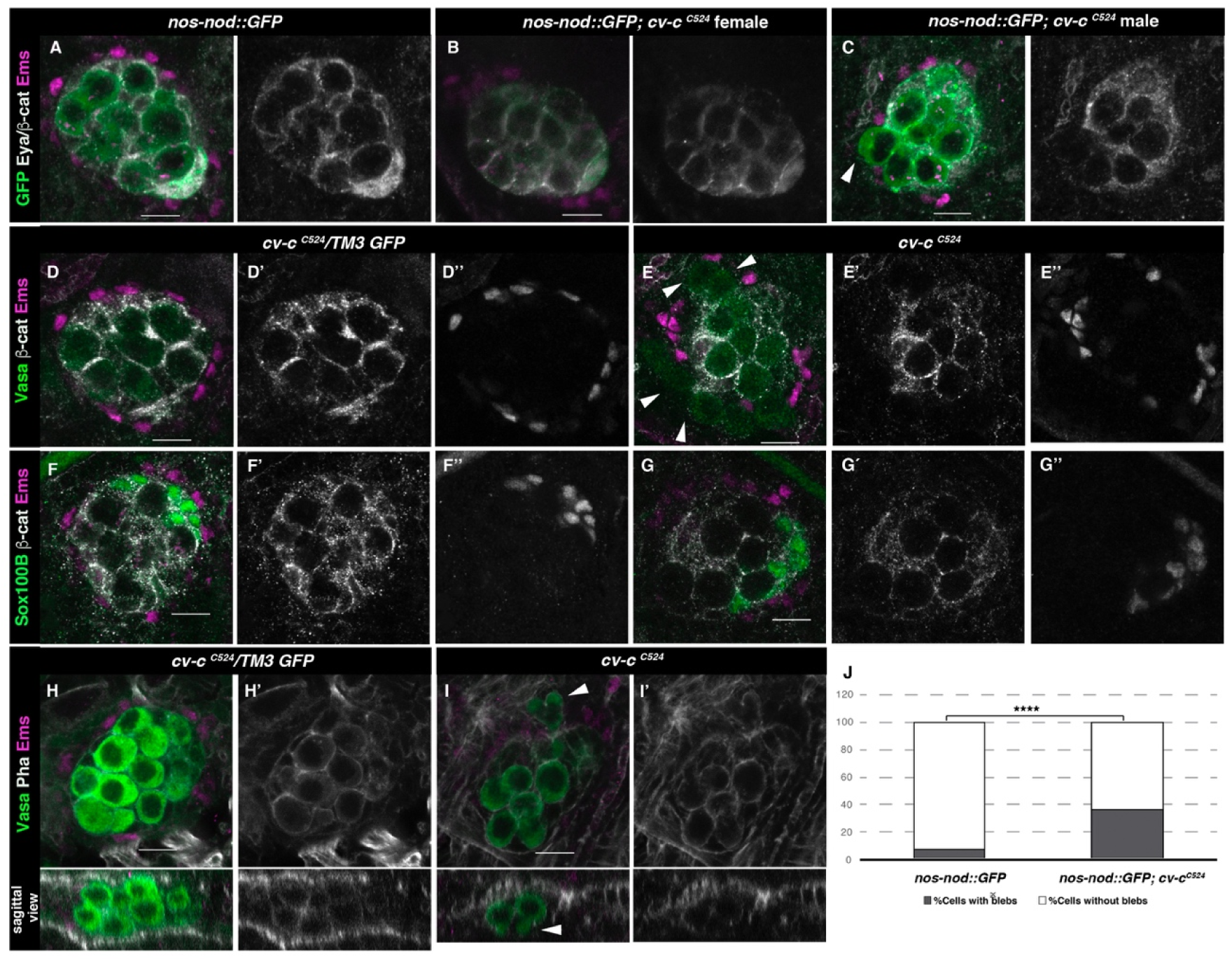
Gonad morphology in *cv-c* mutant embryos. (**A**-**C**) Gonads with germ cells labelled with *nos-nod::GFP* (green), pigment cells with anti-Ems (magenta) and msSGP with anti-Eya (nuclear grey staining) and the AJs with anti-β-catenin (grey membranes). Right panels show Eya and β-catenin channel. (**A**) In the control testis, germ cells are ensheathed by thin mesoderm extensions produced by the interstitial cells detectable by β-catenin staining. Similar germ cell ensheathment is observed in *cv-c^C524^* ovaries (**B**), while in *cv-c^C524^* testis (**C**) some germ cells become extruded from the gonad and are not enveloped by β-catenin (arrowhead). (**D**-**G**) Testes labelled with mesodermal specific markers to detect the pigment cells (Ems, magenta D-G) or the msSGPs (Sox100B, green F-G) in heterozygous (**D**, **F**) or *cv-c^C524^* homozygous mutant embryos (**E**, **G**). Right panels correspond to β-catenin channel (D-G) first; followed by Ems (D, E) or Sox100B (F, G) channels. All mesoderm cell types are specified in *cv-c* mutant testis despite morphological aberrations resulting in the pigment cell layeŕs discontinuity (compare D and F with E and G). (**H**-**J**) Testes stained with anti-Vasa (green) to label the germ cells and Phalloidin (grey and right panels) to show Actin filaments in heterozygous (**H**) or homozygous *cv-c^C524^* mutants (**I**). Germ cells in mutant testes present protrusions compatible with migratory movements (arrowheads). Z sections are shown below H, I panels. Scale bar 10 µm. (**J**) Quantification of blebbing cells in wild-type or mutant cv-c background using Fisher-test; p-value < 2.2e-16.

To test if these defects are due to the abnormal migration of the germ cells or the mesoderm gonadal precursors during early gonad organogenesis [up to gonad coalescence at stage 15 (st15)] or to later defects on testis maintenance, we labelled the germ cells using *nos-nod::GFP* and the mesodermal cells with *six-moe::GFP* to investigate gonad formation *in vivo*. Using this set up we could observe how the migrating germ cells and the somatic cells converge during normal development, coalescing at embryonic st15 to form a stable spherical gonad (Movie 1). Analysis of *cv-c^C524^* homozygous mutant embryos in the same conditions revealed that development is normal up to st15, with the testis compacting into a spherical gonad (Movie 2). However, after this stage the germ cells become extruded from the testis (Movies 2 and 3). Analysis of fixed mutant testes shows that the extruded germ cells extend blebs that are more characteristic of the earlier migratory phase (Fig. 3*I* to J arrowheads) (Jaglarz & Howard, 1995). These blebs are not observed in the ovaries of *cv-c* mutant females (Supplementary Fig. *2B*) nor in the wild type testis after gonad compaction (Fig. 3*H*).

### Rescue of *cv-c Drosophila* testis defects with human DLC3

We have previously shown that DLC3 can rescue the mutant phenotypes caused by *cv-c* mutations in the Malpighian tubules, the kidney-like structures of the fly, indicating these homologous *Drosophila* and human proteins conserve similar functions (Sotillos *et al*., 2018). Therefore, given that Cv-c is expressed and required in the male *Drosophila* gonad, and that DLC3 can functionally substitute for Cv-c in some tissues, we investigated if DLC3 is also capable of rescuing the mutant gonadal defects observed in *cv-c^C524^* homozygous embryos (Fig. 4*A* and *D*). Using the UAS/Gal4 system to express wild type DLC3 protein with the pan mesodermal *twi-Gal4* driver line in otherwise *cv-c^C524^* mutant embryos, we efficiently rescued the testis defects (Fig. 4*A, B* and *E*) pointing out to the conservation of DLC3/Cv-c function in gonadogenesis. In contrast, expression in the same conditions of the *DLC3^S993N^* StART protein present in human patients is not capable of efficiently rescuing the testis phenotype (Fig. 4*A, B* and *F*).

**Figure 4.**
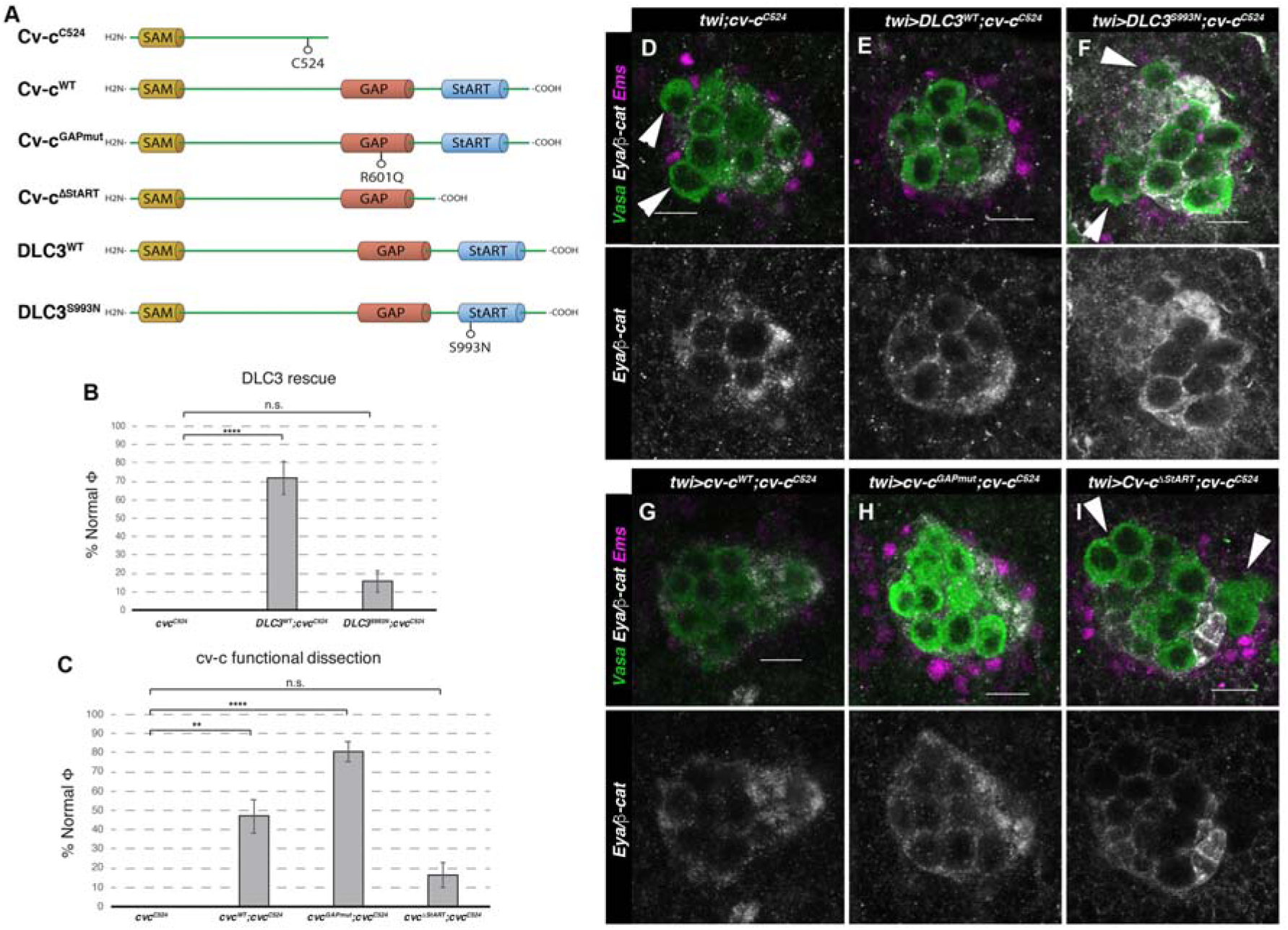
Rescue of *cv-c* mutant testes. (**A**) Schematic representation of Cv-c and DLC3 protein variants studied. (**B**-**C**) Rescue of the dysgenic testis of *cv-c^C524^* homozygous mutant males after expressing the specified (B) DLC3 or (C) Cv-c protein variants under UAS control with the pan mesodermal *twi-Gal4* line. Phenotypic rescue is shown as percentage of normal testes quantified as encapsulated gonads with all germ cells inside the testis as assessed by Fisher-test. Representative images of testes in (**D**) control homozygous *cv-c^C524^* animals, or homozygous *cv-c^C524^* animals expressing in the mesoderm either (**E**) *UAS-DLC3^WT^*, (**F**) *UAS-DLC3^S993N^*, (**G**) *UAS-cv-c^WT^,* (**H**) *UAS-cv-c^GAPmut^* or (**I**) *UAS-cv-c^ΔStART^*. Arrows in D, F and I point to extruded germ cells that are not surrounded by β-catenin. Testes are stained with anti-Vasa to label the germ cells (green), anti-Ems to label the pigment cells (purple) and anti-Eya and anti-β-catenin to label the msSGPs and the membranes ensheathing the germ cells respectively (grey in lower panels). Scale bar: 10 µm Fisher-test; ns, p > 0.05; **p<0.001; ****p < 0.0001.

### Testis development in *Drosophila* requires the StART domain

To elucidate the molecular mechanisms mediating DLC3/Cv-c function in gonad development, we analysed the capacity of different Cv-c protein variants to rescue the testis defects of *cv-c^c524^* homozygous embryos (Fig. 4*A*).

As mentioned above, StART domain mutations in DLC3 are the suspected cause of gonadal dysgenesis in human patients. In agreement with this, we found that expression of a *UAS-cv-c*^ΔStART^ construct generating a Cv–c protein lacking the StART domain, does not significantly normalize the testis defects (Fig. 4*C* and *I*). In comparison, the expression of the wild type Cv-c protein rescued the abnormal phenotypes in more than 50% of the testes (Fig. 4*C* and *G*). Surprisingly, expression of a Cv-c mutant protein substituting a highly conserved Arginine into Glutamine residue that has been shown to block the GAP domain activity *in vitro* and the protein function *in vivo* (Leonard *et al*, 1998; Sotillos *et al*., 2013; Sotillos *et al*., 2018) rescued the gonadal phenotypes to a better extent than the wild type protein (Fig. 4*C* and *H*). This may be due to the overexpression of a functional GAP protein resulting toxic, not allowing to appreciate the full rescue of the StART-mediated function, a phenomenon that has been described previously (Hendrick & Olayioye, 2019; Holeiter *et al*, 2012). This does not happen in Cv-c^GAPmut^ nor in DLC3^WT^, which may have a less efficient GAP function in a *Drosophila* environment than Cv-c.

Moreover, analysis of embryos homozygous for *cv-c*^7^, an allele which carries that exact GAP mutation in the endogenous gene (Denholm *et al*., 2005), showed normal testis (Fig. 5*A* and *B*), suggesting that Cv-c function in the gonad is not mediated through its RhoGAP function, but requires the StART domain, deleted in *cv-c^C524^* (Fig. 5*C* and *D*) and *cv-c^M62^* mutant alleles (Fig. 5*E* and *F*). To test if these results are due to Cv-c function in the testis, we repeated the experiments using *c587-Gal4,* a line expressed specifically in the gonad mesoderm cells ensheathing the germ cells. Although with a lower penetrance, this line results in similar rescues indicating a specific requirement of Cv-c in the testis gonadal mesoderm (Supplementary Fig. *S3*).

**Figure 5.**
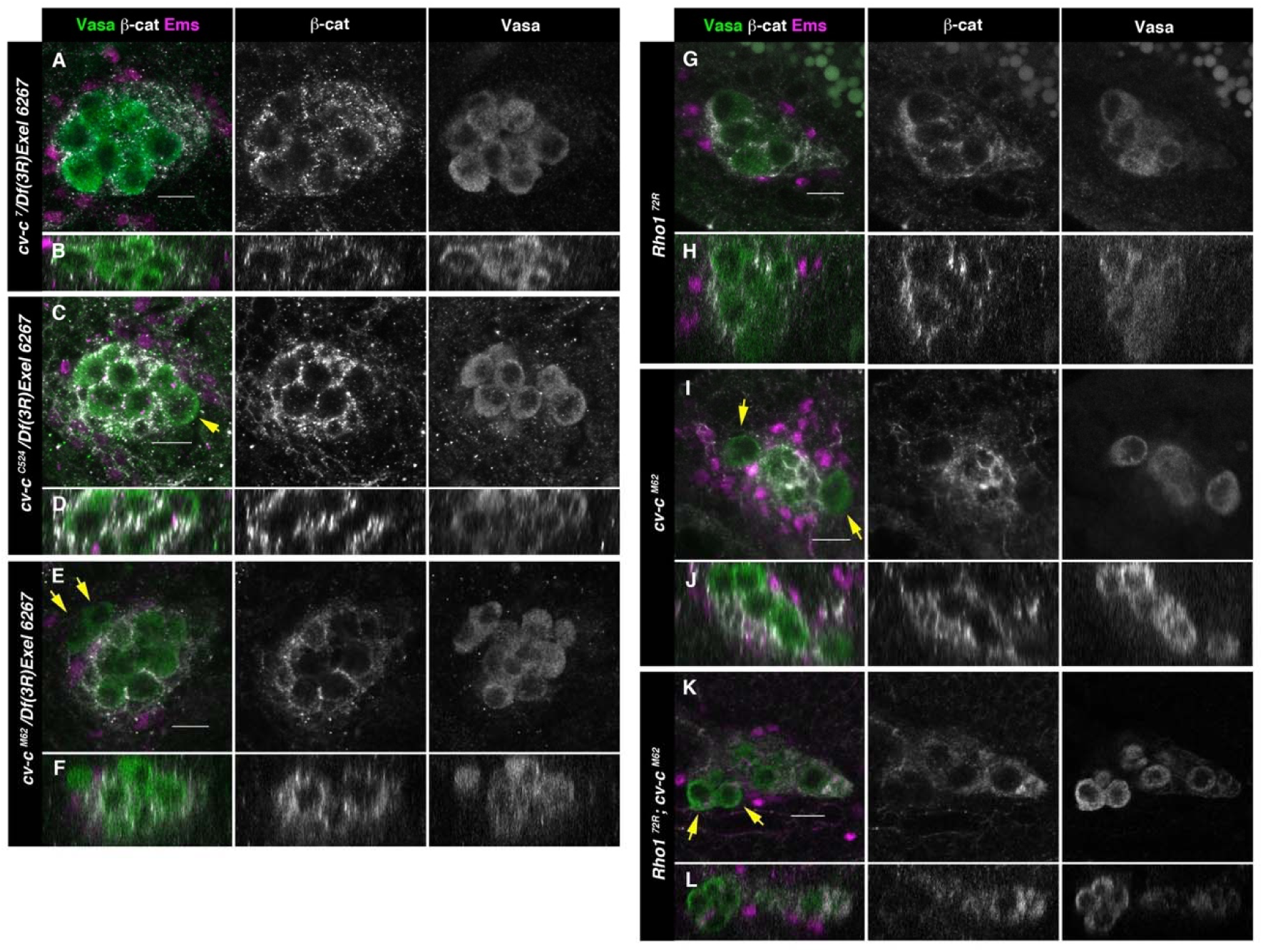
Testes in embryos with altered small GTPase regulation. (A-L) Testes of various genotypes stained with anti-Vasa (green), β-catenin (grey) and Ems (magenta). β-catenin and Vasa channels are shown separately in right panels. (**A**, **B**) No extruded gem cells are observed in hemizygous *cv-c^7^*/*Df(3R)cv-c Exel6267* embryos carrying a *cv-c^7^* allele which inactivates the RhoGAP domain’s function. (**C**-**D**) Hemizygous *cv-c^C524^*/*Df(3R)cv-c Exel6267* or (**E**-**F**) *cv-c^M62^*/*Df(3R)cv-c Exel6267* show extruded germ cells. (**G**-**H**) *Rho1^72R^*mutant embryos have smaller testes without extruded germ cells. (**I**-**J**) Homozygous *cv-c^M62^* mutants present extruded germ cells. (**K**-**L**) Homozygous *Rho1^72R^ cv-c^M62^* double mutant embryos present smaller testis with extruded germ cells. Arrows point to GCs outside the gonads. Note that β-catenin envelops all germ cells in (A-B) and (G-H) while in (C-E) and (I-L) some germ cells are not surrounded (arrows). Z sections are shown under all panels. Scale bar 10 µm.

Although DLC3 and Cv-c are RhoGAP proteins that when mutant cause Rho1 over-activation, our experiments suggest that in the gonad these proteins do not require the GAP function. To confirm that the testis phenotype is not due to Rho1 over-activation we analysed if *cv-c^M62^* gonad mutant phenotypes can be rescued by a *Rho1* mutation. As previously described, in *Rho1* homozygous mutants the gonad precursor germ cell migration is less efficient, giving rise to smaller male and female gonads (Kunwar *et al*, 2003). However, in *Rho1* mutants, the cells reaching the gonads become ensheathed and coalesce to form stable testes as in the wild type (Fig. 5*G* and *H*). In *Rho1 cv-c* double mutant embryos we observe an additive effect of both mutations (Fig. 5*K* and *L*), with smaller gonads due to the *Rho1* early migratory defect and the testis germ cell extrusion typical of *cv-c* mutants (Fig. 5*I* and *J*), indicating that the dysgenic gonad phenotype cannot be rescued by a reduction of Rho1 function.

These results show that the StART domain function is required for human and *Drosophila* testis formation strongly supporting that the dysgenic gonad defects observed in patients are caused by the DLC3 StART mutation and demonstrating that Cv-c has a GAP-independent function that requires the StART domain.

### Cellular causes for testis dysgenesis

We next searched for the underlying cellular defects responsible for the observed testis dysgenesis in *cv-c* mutants. To explore if the germ cell extrusion is due to a failure of the somatic gonadal cells to ensheath the germ cells, we studied the expression of E-cadherin (E-cad), which localizes to the membrane of both germ cells and somatic mesodermal cells and is required for germ cell ensheathment and gonadal compaction (Jenkins *et al*, 2003). We observe that while, in the wild type testes the germ cells inside the gonad are encircled by high levels of E-cad protein (Fig. 6*A* and *A’*), in *cv-c* mutants there are frequent gaps of E-cad distribution between adjacent germ cells inside the testes. In addition, we observe that extruded germ cells have almost no E-Cad on their membranes (Fig. 6*B* and *B’* arrowheads). We observe analogous, abnormal β-catenin (Fig. 3) localisation inside the testes, indicating that the relationship between the germ cells and the surrounding somatic cells is not well established or poorly sustained (Fig. 3*C, E* and *G*).

**Figure 6.**
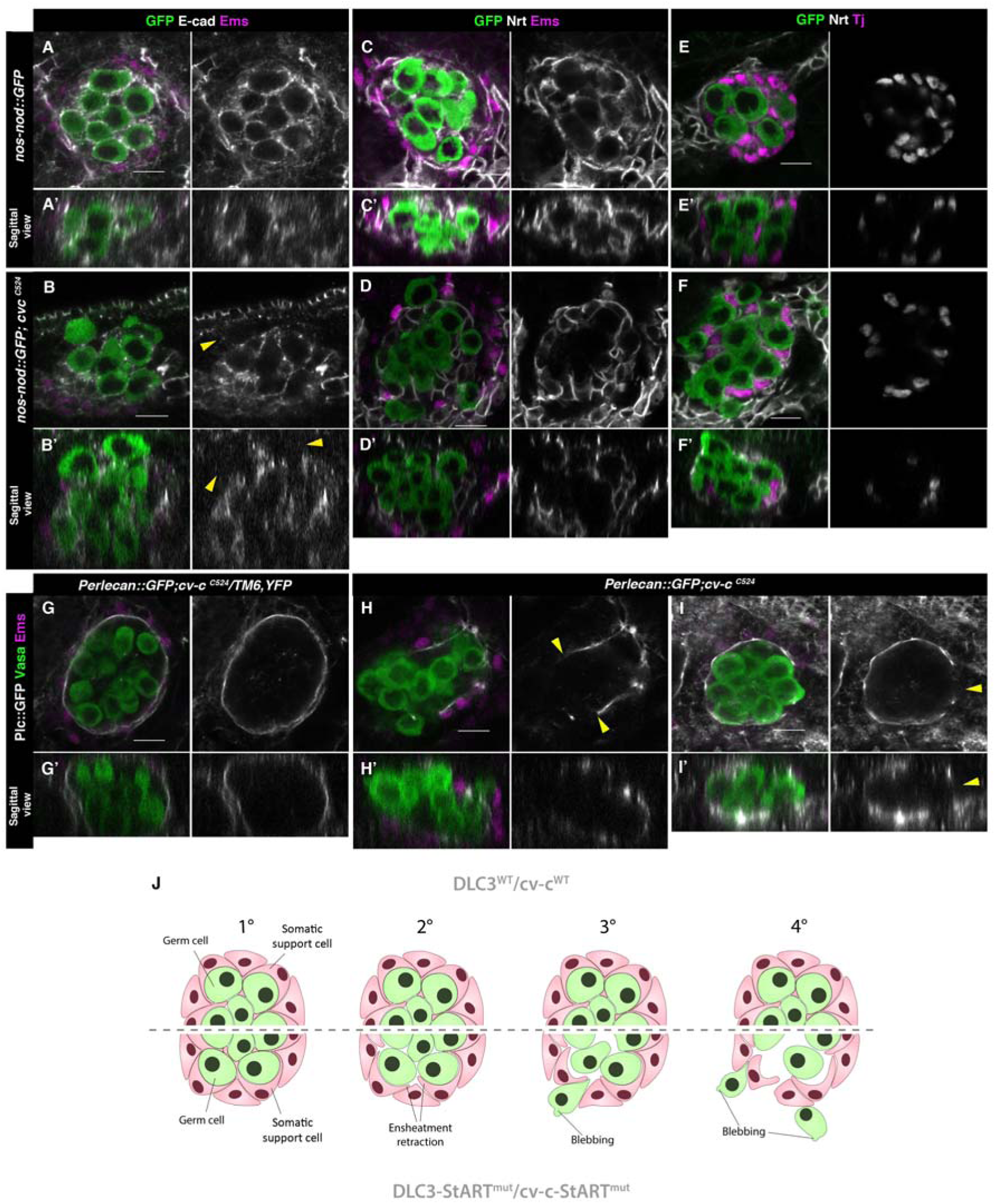
Ensheathment defects in *cv-c* mutant testes. (**A**, **C**, **E**) Wild type and (**B**, **D**, **F**) *cv-c^C524^*testes. Germ cells are labelled in green with *nos-nod::GFP* (green A to F) and in grey with anti-E-cad (A to B) or anti-Nrt (C to D) to highlight the ensheathing membranes (grey in right panels). (**A**, **A’**) In the wild type E-cad highlights contacts between the germ cells and the interstitial somatic gonadal cells surrounding each germ cell reflecting their correct ensheathment. (**B**, **B’**) In *cv-c* mutant testes several germ cells inside the testis and all the extruded ones (arrowheads) are not surrounded by E-cad labelling membranes indicating incorrect ensheathment. (**C**, **C’**) Neurotactin in the wild type testis reflects correct germ cell ensheathment. (**D**, **D’**) In *cv-c^C524^* mutants little Neurotactin expression is detected inside the testis. Nuclei of Tj labelled somatic gonadal cells (magenta and grey in right panels) are detected between the GCs inside wild type testes (**E**, **E’**) but not in *cv-c^C524^*mutant testes (**F**, **F’**). (**G, G’**) Heterozygous and (**H to I’**) homozygous *cv-c^C524^* testes labelling the extracellular matrix with Perlecan::GFP (Plc, grey in right panels) and stained with anti-Vasa (green) and anti-Ems (magenta) to label the GCs and the pigment cells respectively. (G and G’) The wild type testis is enclosed by a Perlecan containing extracellular matrix (green). In *cv-c^C524^* mutant testis a discontinuous extracellular matrix is observed where the GCs are outside the gonad (H to I’, yellow arrowheads). Z sections are shown under all panels. Scale bar: 10 µm. (**J**) Interpretation of the testis degeneration in wild type and DLC3/Cv-c mutants. (1) The StART domain of DLC3/Cv-c has a Rho-independent function stabilizing cell interactions between germ cells and somatic support cells. Stable cell-cell interaction among the cells of the gonadal niche allows them to settle down in the testis. (2) In DLC3^S997N^/Cv-c^ΔStART^ testis cell-cell interactions become compromised, cells lose their cohesion and separate from the gonadal niche. (3) After losing communication with somatic cells, germ cells become extruded from the gonadal niche initiating an erratic migrating behaviour. (4) Progressive loss of germ cells leads to gonad degeneration.

In addition to E-Cad, we also analysed Neurotactin (Nrt) expression that, in normal fly embryos, localises to the cell extensions produced by the interstitial mesodermal cells that ensheath the germ cells (Fig. 6*C* and *C’*) (Jenkins *et al*., 2003). In *cv-c* mutant testes we find that Neurotactin expression is almost absent around the germ cells inside the gonad (Fig. 6*D* and *D’*). Accordingly, interstitial somatic gonad mesodermal cells (labelled by the Traffic jam (Tj) antibody), which in the wild type gonad can be detected distributed between GCs (Fig. 6*E* and *E’*), were frequently displaced to the periphery in *cv-c* mutant testes (Fig. 6*F* and *F’*).

Finally, we studied the gonad integrity in *cv-c* mutants. Wild type testes are surrounded by a Perlecan rich extracellular matrix located between the pigment cells and the interstitial mesodermal cells that can be detected using a GFP insertion in the *Perlecan* gene (Fig. 6*G* and *G’*). In *cv-c^C524^*mutants, the pigment cell layer and the ECM matrix are discontinuous with the extruded germ cells locating where the matrix gaps appeared (Fig. 6*H* to *I’*, arrowheads). These findings suggest that germ cell ensheathment by somatic cells cannot be maintained in the testis of *cv-c* mutant flies resulting in germ cell extrusion and gonadal rupture, a phenotype that is rescued by both Cv-c and DLC3 proteins containing a wild type StART domain. Taken together our results show that the DLC3/Cv-c protein family has a StART dependent function required for male gonadogenesis in humans and *Drosophila*.

## Discussion

Sex development is a central event in the life of metazoan animals. Sexual reproduction depends, particularly in mammals, on the determination of the gonads. What makes the gonads unique among all other organs is the fact that from one primordium two morphologically and functionally distinct organs, a testis and an ovary, arise. This phenomenon requires the fine tuning in space and time of all the genes, cells and processes involved (Carre & Greenfield, 2016). Although new players are continuously discovered, many mechanisms underlying sexual determination and differentiation remain poorly understood, often due to the lack of a reliable experimental model. With this work, using *Drosophila* as a model, we confirmed the implication of *DLC3* as a novel DSD gene required for testis determination through the action of its StART domain.

*DLC3* had been found to be mutated in two 46, XY DSD siblings presenting gonadal dysgenesis but no experimental evidence confirming causality had been provided (Ilaslan *et al*., 2018). In this study we report a third DSD patient with a mutation in DLC supporting the involvement of this gene in gonad development. First described in human myeloid cells, *DLC3* loss of expression was found in primary tumours from different tissues (Durkin *et al*, 2007). This multidomain protein forms part of a RhoGAP conserved family containing an N-terminus SAM domain followed by a serine-rich region, a catalytic GAP domain, and a StART domain (reviewed in (Braun & Olayioye, 2015)). There are also alternative isoforms lacking the SAM domain such as DLC3-Δ (Durkin *et al*., 2007). Despite the recent advances identifying its structure and spatial subcellular location (Braun *et al*, 2015; Durkin *et al*., 2007; Hendrick *et al*, 2016; Holeiter *et al*., 2012), the specific function of the various protein domains is still not completely understood. At the adherent junctions (AJ) of epithelial cells, DLC3-GAP domain inhibitory effect on RhoA stabilizes E-cadherin-based cell–cell contacts (Hendrick & Olayioye, 2019). DLC3 is also known to influence the dynamics of the AJ in an indirect way, regulating E-cadherin turnover at the recycling compartments (Braun *et al*., 2015).

In search for the molecular elements mediating DLC3/Cv-c testis function, we studied the involvement of the StART and GAP domains. In the fly, Cv-c is expressed in the testis mesodermal cells during development and *cv-c* mutations deleting the GAP and the StART domains disrupt testis development without affecting ovarian development. Expression of the DLC3 human protein in *Drosophila* mutants can substantially rescue the male gonadal defects while the expression of the mutant DLC3^S993N^ form found in DSD patients does not, proving a conserved functionality among both species and the implication of DLC3 as a novel DSD gene.

The only enzymatically active domain previously recognised in DLC3/Cv-c was the GAP domain with the StART domain thought to play a regulatory role. Surprisingly, the expression of Cv-c without a GAP-functional domain in a *cv-c* mutant background is able to rescue the testis’ developmental disruption with a similar efficiency than wild type DLC3 transgenic flies. However, transgenes without the StART domain (Cv-c^ΔStART^) or the human DLC3^S993N^ allele carrying a mutation in the StART domain were unable to rescue the male gonad defects. These results demonstrate that the Cv-c StART domain has a GAP-independent function required for male gonad formation in *Drosophila*. StART domains have been shown to be lipid binding (Gatta *et al*., 2018; Horenkamp *et al*., 2018; Iaea *et al*., 2015; Khelashvili *et al*., 2019; Naito *et al*., 2019). The observation of a disrupted somatic-germ cell niche in the testis of *cv-c* mutant flies, made us wonder whether the StART domain’s function in gonad development might be linked to lipid-regulated cell-cell interactions.

Our modelling analysis predicts that the S993N DLC3 mutation affects the Ω1-loop structure of the StART domain. Ω-loops play multiple roles in protein function, often related to ligand binding, stability, and folding (Fetrow, 1995). This loop is conserved in the StART domains of several other STARD/DLC proteins and seems to be functionally important to modulate access to the ligand-binding cavity (Gatta *et al*., 2018; Gatta *et al*, 2015; Horenkamp *et al*., 2018; Naito *et al*., 2019). Our *in-silico* analysis of the DLC3-StART domain predicts the Ω1-loop displays the highest frequency of interaction with the membrane. In line with these, *in silico* simulation of the DLC3^S993N^ mutant StART domain predicted the loss of one of the two conformational states of the protein, impairing its conformational dynamic with the loss of a close state ligand-binding pocket in the StART domain. Within the loop, we find that residues S993, M994, A915, P996 and H997, have the highest frequency of interaction with the lipid bilayer. Interestingly, these five residues are conserved in the StART domains of StARD12 and StARD13, with the proline residue (P996 in DLC3) also conserved in StARD2, StARD7, StARD10 and StARD11 (Thorsell *et al*, 2011). These results suggest that the mutated S993N residue affects a critical structure of the protein domain and that alterations of this Ω1-loop could impair the domain’s membrane binding activity independently from the GAP domain. In this context, single mutations in the Ω1-loop surroundings, like the one carried by the patients, may have drastic effects on overall protein stability with consequences on the maintenance of the junctions between gonadal precursor cells.

In vertebrates and flies, gonad formation requires that germ cells migrate to the gonadal microenvironment, interact with the somatic cells, and cease the migrating behavior. Germ cells migrate following an increasing gradient of chemo-attracting signals (Richardson & Lehmann, 2010). These signals trigger cell polarization in germ cells, reorganizing the membrane’s cadherins. The protein rearrangement is followed by a reorganization of the cytoskeleton, creating a pulsation that flows front to back (Kardash *et al*, 2010; Kunwar *et al*, 2008; Kunwar *et al*., 2003). During this process there is a characteristic formation of “blebs” or protrusions at the front of the migrating cell (Kardash *et al*., 2010). Germ cells migrate until they reach a point of maximum chemoattractant concentration. However, when subjected to ectopic signals, germ cells continue to show protrusions and migrating behavior without a clear polarization (Kardash *et al*., 2010; Richardson & Lehmann, 2010). The PGCs’ close association with the SGPs end the migratory phase, the last PGCs divisions are detected prior to compaction completion and the germ cells stop extending protrusions entering an “inactive” phase (Jaglarz & Howard, 1995).

We show that in the absence of Cv-c function in the *Drosophila* testis, the mesodermal pigment cells do not form a continuous layer around the testis and the ECM surrounding the testis breaks. We also show that the interstitial gonadal cells fail to ensheath the germ cells and as a result of all these the germ cells become extruded from the testis. These perturbations can be partially corrected by expression in the testis mesoderm of human DLC3 or *Drosophila* Cv-c that in both cases require a functional StART domain. Thus, our results suggest that Cv-c/DLC3 have a fundamental function on the mesodermal testis cells that has been conserved. These results indicate that, as in *Drosophila*, the primary cause for the gonadal dysgenesis in DLC3 human patients is due to the abnormal maintenance of the testis mesoderm cells, which include both Sertoli and Leydig cells.

The germ and somatic gonadal cells interact via the formation of AJs in sex-specific patterns (Fleming *et al*, 2012). The junction protein complex in male gonads is relatively well conserved among species, with its core constituted by head-to-head cadherin dimers established from opposing interacting cells (Piprek *et al*, 2020; Troyanovsky, 2012). As in *Drosophila*, human PGCs have to migrate from the allantois to the genital ridge where the gonad is formed and later they interact with the Sertoli mesodermal cells. This interaction is essential for the PGCs to differentiate and survive and requires, both in humans and *Drosophila*, an E-cad levels increase when the PGCs and SGPs meet (Fleming *et al*., 2012; Kunwar *et al*., 2008). This initial interaction is fragile and must be stabilized by incorporating β-catenin and other proteins to the complex (Fleming *et al*., 2012; Piprek *et al*., 2020). The complex serves finally as an anchor for the actin filaments of the cytoskeleton. Accordingly, the alteration of E-cadherin and β-catenin distribution observed in *cv-c* and *DLC3* (humanized) fly mutant male gonads, together with the appearance of “blebs”, suggest that the settling-down switch has not been activated in the affected cells, and they remain in an erratic migrating behavior until they eventually escape from the gonadal niche (summarized in Fig. 8*J*).

During the compaction stage, the *Drosophila* somatic cell’s E-cad locates into thin membrane extensions that surround each germ cell, which also expresses E-cad (Jenkins *et al*., 2003). In our study, we observed that E-cadherin and β-catenin were abnormally distributed in *cv-c* mutants, suggesting alterations of the connectivity between PGCs and SGPs. Interestingly, this phenomenon was observed even under Rho1 reduction, reinforcing the idea of a role of Cv-c in a Rho1-independent mechanism that promotes the stabilization of AJ after gonad coalescence.

This Rho-independent mechanism requires StART domain’s integrity. Although most of the previous efforts to explain the molecular function of DLC3-StART domain have failed (Alpy & Tomasetto, 2005), Beatriz Sanchez-Solana et al. (Sanchez-Solana *et al*, 2021) proved that the DLC1-3 StART domains bind phosphatidylserine (PS). They also postulated that, in DLC1, the lipid-binding works as a mediator of the interaction with several proteins, independently from Rho-GAP domain activity (Sanchez-Solana *et al*., 2021). In eukaryotic membranes, PS is a well-known phospholipid involved in signalling pathways (Kay & Grinstein, 2013). In healthy cells, most PS locates on the inner layer of the plasma membrane. When the AJs are established, PS forms trans-bilayer-coupled nanoclusters with GPI-anchored proteins and glycosphingolipids from the outer layer. PS microdomains become anchoring points for proteins that promote signal transduction and the stabilization of the junction like Flotillins and PS-binding proteins (Yap *et al*, 2015). Taking these findings together, we hypothesise a role for the StART domain in the interaction of DLC3 with the temporal PS nanoclusters formed at the AJs, a phenomenon already demonstrated for DLC1 (Sanchez-Solana *et al*., 2021). Mutated DLC3-StART domain lacking the ability to respond to lipid-binding with a conformational change, could be incapacitated to promote any further signalling.

The coincident requirement of DLC3/Cv-c for testis development and the conservation of the StART function suggest that in humans the DLC3-StART domain activity could also be required for the Sertoli cells/SGPs to trigger the germ cell settling behavior and consolidate the male gonad development. It is still unknown why ovaries, where a similar somatic ensheathment of the germ cells occurs, do not require DLC3/Cv-c to maintain its stability. A possible explanation could be the sex-specific influence of AJ in overall patterning of the testis vs. the ovary at the time of early gonadal sex differentiation (Fleming *et al*., 2012).

In conclusion, we demonstrate that mutations in DLC3/Cv-c are a novel cause of testicular dysgenesis in *Drosophila* and humans. Our results suggest that the dysgenesis is caused by the observed destabilization of cell-cell connexions between testis mesodermal cells and between the germ cells and somatic support cells after gonad coalescence. DLC3/Cv-c action in the gonads sheds new lights on the mechanism by which the germ cells end their migrating behaviour and settle in the gonadal niche. Pending the analysis of DLC3/Cv-c in further species, our results indicate this function could be a conserved mechanism among species, since human DLC3 is able to rescue the *cv-c* testicular defects in *Drosophila* embryos. Our work points to DLC3/Cv-c as a novel gene required specifically in testis formation. Adding DLC3 to the list of genes involved in 46X,Y complete dysgenesis opens up a new avenue to analyse the molecular and cellular mechanisms behind these disorders that could help in diagnosis and the development of future treatments. These results also underline the outstanding potential of *Drosophila* as a model to unveil the functional mechanism underlying human conditions like DSD without the ethical and logistical complexity of more conventional mammalian models.

## Materials and Methods

### Molecular Dynamics Simulations

The structure of the StART domain of human DLC3/StARD8 (Uniprot ID Q92502) was obtained from Alphafold (AF) (Jumper *et al*, 2021). A truncated structure of the domain comprising residues 838 to 1012 was used. The WT and S993N systems were set up using the CHARMM-GUI Solution Builder (Jo *et al*, 2008) with a cubic box of edge length of 7.3 nm. The systems were solvated with TIP3P water and ionized with 0.12M of sodium and chloride ions. 3 independent replicas of 100 ns each were simulated for each system using the GROMACS (Van Der Spoel *et al*, 2005) 2018.6 package and the CHARMM36m force field (Huang *et al*, 2017). Initial equilibration was carried out by performing energy minimization using the steepest descent algorithm, followed by a short NVT and NPT equilibration of 100 ps each with position restraints on the backbone atoms of the protein. Production runs were performed at 310K using a velocity-rescale thermostat (Bussi *et al*, 2007), with separate temperature coupling for protein and solvent particles, and a coupling time constant of 0.1 ps. The first 10 ns of the production runs were not considered for analysis. The md integrator was used for the production runs, with a time step of 2 fs. The Parrinello-Rahman barostat (Parrinello & Rahman, 1981) was used to maintain the pressure at 1 bar, with an isotropic pressure coupling scheme, a compressibility of 4.5 x 10-5 bar-1 and a coupling time constant of 2.0 ps. Electrostatic interactions were evaluated using Particle Mesh Ewald (PME) with a Fourier spacing of 0.16 nm, a cutoff of 1.2 nm, and an interpolation order of 4. _Van der Waals (VDW) interactions were switched to zero over 10 to 12 Å. Bonds involving hydrogen atoms were constrained using the LINCS algorithm Periodic boundary conditions were used in all three dimensions.

The distance between the Cα atoms of the residues were computed every 100 ps and the distribution was obtained using Kernel Density Estimation (KDE). Fig. 1*C* was rendered using Visual Molecular Dynamics (VMD) (Humphrey *et al*, 1996).

Eight independent replicas of 3 µs each were simulated using the GROMACS (2018.x) (Van Der Spoel *et al*., 2005) package and the Martini 3 force field61. Initial equilibration was carried out by performing energy minimization using the steepest descent algorithm, followed by a short MD run of 250 ps. Production runs were performed at 310K using a velocity-rescale thermostat (Bussi *et al*., 2007), with separate temperature coupling for protein, bilayer, and solvent particles and a coupling time constant of 1.0 ps. The md integrator was used for the production runs, with a time step of 20 fs. The Parrinello-Rahman barostat (Parrinello & Rahman, 1981) was used to maintain the pressure at 1 bar, with a semi-isotropic pressure coupling scheme and a coupling time constant of 12.0 ps. The Coulombic terms were calculated using reaction-field (Tironi *et al*, 1995) and a cut-off distance of 1.1 nm. A cutoff scheme was used for the vdW terms, with a cut-off distance of 1.1 nm and Verlet cut-off scheme for the potential-shift (Verlet, 1967). The Verlet neighbour search algorithm (Páll & Hess, 2013) was used to update the neighbour list every 20 steps with a buffer tolerance of 0.005 kJ mol−1 ps−1. Periodic boundary conditions were used in all three dimensions. The system setup and simulation parameters are in line with the recently proposed protocol for studying transient protein-membrane interactions with the Martini force field (Srinivasan *et al*, 2021). Membrane binding events were assessed using the time-trace of the minimum distance between the protein and the membrane, computed with the gmx mindist tool in GROMACS. Membrane-interacting residues were computed every 500 ps using an in-house script with the following protocol: a residue was considered to interact with the membrane if the distance between any bead of the residue and any lipid bead was lower than or equal to 0.5 nm. For each residue, we counted the instances of residue-membrane interaction during the trajectory, summed this value over all the replicas, and computed a corresponding normalized value. Figures. 2A and 2D were rendered using Visual Molecular Dynamics (VMD) (Humphrey *et al*., 1996).

### Fly strains

The following lines: *Mi{PT-GFSTF.0}cv-cMI00245-GFSTF.0*, *Sxl::GFP* and *Df(3R)Exel 6267* were obtained from the Bloomington Drosophila Stock Center. The Cv-c::GFP fusion protein results from the integration into cv-c of a gfp sequence flanked by acceptor/donor splicing sequences present in a transgene (*Mi{MIC}t cv-c MI00245*) inserted into the third intron of the endogenous gene (Venken *et al*, 2011). *cv-c^M62^*, *cv-c ^C524^*, *cv-c^7^* were described in Denholm et al. (Denholm *et al*., 2005). *UAS-cv-c^WT^*, *UAS-cv-c^GAPmut^*, *UAS-cv-c^ΔStART^* and *UAS-Myc-DLC3^WT^* were described in (Sotillos *et al*., 2018). *nos-nod::GFP* (is a gift from A. González Reyes). *c587-Gal4* (is a gift from E. Matunis). *six4-moe::GFP* (Sano *et al*, 2012) and *Vasa::GFP* (a gift from P. Lasko). *Perlecan::GFP* was described in (Morin *et al*, 2001). *Rho1^72R^* was described in (Strutt *et al*, 1997). *twist-G4* was described in (Greig & Akam, 1993).

Rescue experiments were performed crossing *twist-Gal4/CyOwg-lacZ; cv-c^C524^/TM6B risk-lacZ* females with *UAS-X/CyOwg-lacZ; cv-c^C524^/TM6B risk-lacZ* males or *c587-Gal4; cv-cC524/TM6B risk-lacZ* females with *UAS-X/CyOwg-lacZ; cv-c^C524^/TM6B risk-lacZ* males (where X corresponds to either *cv-c^WT^*; *cv-c^GAPmut^*; *cv-c^ΔStART^*; *Myc-DLC3^WT^*or *Myc-DLC3^S993N^*).

Flies were raised at 25°C.

### Constructs

To generate *UAS-Myc-DLC3^S993N^* a XhoI DLC3^WT^ fragment was subcloned into pBlueScript (DLC3-Xho-pBs). DLC3-Xho-pBs was used as a template to mutate Ser at position 993 to Asn of human DLC3 (CCDS48134.1) using Pfu Polymerase.

The following primers were used:

Forward: 5’-TGTACCACTATGTCACCGACA-A-CATGGCACC-3’

Reverse: 5’-TGGGGTGCCATG-T-TGTCGGTGACATAGTG-3’

After PCR reaction DNA was incubated with DpnI during an hour at 37° to digest the methylated template DNA and transformed. Clones were sequenced by standard methods.

From the selected clone (DLC3^S993N^-Xho-pBs) a BglII fragment containing the mutation was substituted in the pUASt-Myc-DLC3^WT^ to obtain pUASt-Myc-DLC3^S993N^.

Constructs were injected by the *Drosophila* Consolider-Ingenio 2007 transformation platform (Spain).

### Immunohistochemistry

Embryos were collected on apple juice agar plates that contained yeast paste. For inmunostaining experiments female flies were allowed to lay eggs overnight onto plates at 25°C. Embryos of an overnight lay were dechorionated 2’ in bleach dilution in water (1:1) and fixed for 20 minutes in a PBS-formaldehyde 4%/heptane mix. After removing the fixative, methanol was added and shaken vigorously for one minute to remove the vitelline membrane. After allowing both phases separate, sinking embryos were recovered washed in clean methanol and rehydrated in PBS-Tween 0.1% (PBT) and preabsorbed during an hour in PBT-1%BSA at room temperature (RT). Primary antibodies were used at the described concentration, diluted in PBT-1%BSA and incubated overnight at 4°C. Primary antibodies were washed 2X5 minutes in PBT and preabsorbed 1h in PBT-1%BSA at RT. Embryos were incubated with the secondary antibody diluted at 1:400 in PBT-1%BSA at RT during 4 hours. After incubation embryos were washed 4X15’ in PBT-1%BSA at RT follow to 2 rinses in PBS before mounting in Vectashield.

The following primary antibodies from the Developmental Studies Hybridoma Bank (DSHB) were used: rat anti-Vasa 1:20 (DSHB, VASA), mouse anti-Nrt 1:100 (DSHB, BP106), mouse anti-β-catenin 1:50 (DSHB, N27A1), rat anti-DE-cad 1:50 (DSHB, DCAD2), mouse anti-Eya 1:100 (DSHB, Eya), guinea pig anti-Ems 1:5.000 (gift from U. Walldorf), rabbit-anti-Sox100B 1:1.000 (gift from S. Russell), guinea pig anti-Tj 1:1000 (Gunawan *et al*, 2013), mouse anti-βgal 1:1.000 (Promega, Z378A) rabbit anti-GFP 1:300 (Invitrogen, A11122), chicken anti-GFP 1:500 (Abcam, ab13970).

Secondary antibodies were coupled to Alexa488, Alexa555 or Alexa647 (Molecular Probes). Filamentous Actin was stained with Phalloidin-Rhodamine (Molecular Probes, R415).

Fluorescent in situ hybridization was performed according to standard protocols adding a secondary fluorescent antibody (anti-goat Alexa555, Invitrogen A-21432); cv-c riboprobe was marked using DIG RNA Labelling Kit (Roche, 11 175 025 910). Images were taken on an SPE Leica confocal microscope and processed using FIJI and Adobe Photoshop programs.

### In vivo gonad analysis

In the nos-nod::GFP construct a nanos enhancer drives expression in the germ cells of a microtubule binding GFP protein due to its fusion to the Nod protein fragment. In P-Dsix4-eGFP::Moesin construct a six enhancer drives expression in the male specific somatic cells of the Moesin actin-binding domain fused to GFP (gift from S. DiNardo (58)).

Embryos were dechorionated in bleach and positioned dorsally on top of a coverslip thinly coated with heptane glue and covered with a drop of halocarbon oil. Embryos were imaged on an SP5 Leica confocal microscope, using a 40X oil immersion objective. For each movie, twenty-six to thirty time points were collected. For each time point, between 20 and 40 Z sections were collected (spaced between 0.5 µm and 1 µm). Movies were assembled using IMARIS and Fiji ImageJ software.

### Statistical analysis

Data were obtained from at least 3 biological replicas. Replica samples for each genotype were collected in parallel on different days. Stainings with anti-Vasa, anti-E-Cad and anti-Ems were carried-out to distinguish the GCs, the gonad contour and the testis respectively. We captured confocal z-stacks (at 0.5 µM intervals) encompassing the entire width of testes (typically 40-45 slices). A testis was considered abnormal if one or more germ cells were partially or totally outside the gonad. Data were analyzed by Microsoft Excel and GraphPad Prism. Statistical analyses were performed using Fisher-test and error standard error as described in (Xu *et al*, 2010). Statistical significance was assumed by P<0.05.

## Acknowledgments

This work was supported by a María de Maeztu Unit excellence grants MDM-2016-0687 and CEX-2020-001088-M and a Ministerio de Ciencia e Innovación grant PID2019-104656GB-I00 cofunded by the European Regional Development Fund (FEDER) to J.C.-G.H.

S.V. acknowledges support from the Swiss National Science Foundation (PP00P3_194807). This work was also supported by grants from the Swiss National Supercomputing Centre under project ID s1132. Also, it has received funding from the European Research Council under the European Union’s Horizon 2020 research and innovation program (grant agreement no. 803952).

We thank Acaimo González-Reyes for the critical reading of the manuscript and Steve DiNardo, Paul Lasko, Steve Russell, Uwe Walldorf, Acaimo González-Reyes and Erika Matunis for generously sharing reagents. We also thank the Bloomington Stock Center and the Developmental Studies Hybridoma Bank.

## Ethical statement

All clinical investigations were performed according to the declaration of Helsinki principles. The study was approved by the Geneva ethical committee CCER, authorization number 14-121. The patients and/or their legal guardians gave informed written consent to the study.

---

**Supporting video 1**

*In vivo* gonad coalescence of a control heterozygous *cv-c^C524^/+* testis. The germ cells are labelled with *nos-nod::GFP* and the male specific somatic gonadal precursors with *six-moe::GFP*. Movie taken from st14 (prior to gonad coalescence) up to st16 (after coalescence).

**Supporting video 2**

*In vivo* gonad coalescence of a homozygous *cv-c^C524^* testis. The germ cells are labelled with *nos-nod::GFP* and the male specific somatic gonadal precursors with *six-moe::GFP*. Movie runs from st14 (prior to coalescence) to st16 (after coalescence) when germ cell extrusion becomes noticeable (arrow). Note that the early stages of testis development are normal until the germ cell extrusion begins at later stages.

**Supporting video 3**

Selected planes from the testis presented in Movie 2 to show more clearly the extrusion of an internal germ cell (asterisk).

